# Getting to Know You: Neural Representations of Other People Grow More Perceiver-Specific Over Time

**DOI:** 10.1101/2025.09.23.677973

**Authors:** João F. Guassi Moreira, Kiho Sung, Yixuan Lisa Shen, Sunhae Sul, Yoosik Youm, Carolyn Parkinson

## Abstract

Mental representations of others are central to social behavior, yet little is known about how these representations evolve over significant periods of time as people get to know one another. In this longitudinal functional magnetic resonance imaging study, we tracked how neural representations of familiar peers changed over the course of a school year among two cohorts of first-year high school students (*N* = 150) embedded in newly formed social networks. We assessed the extent to which individuals’ neural representations of their classmates converged with group-level norms—capturing the balance between information that is idiosyncratic to each perceiver and that which is shared across perceivers—and how this changed over time. We identified brain regions where neural representations of familiar peers were well aligned across perceivers, including several areas implicated in person perception, social cognition, and visual processing. Across both cohorts of participants, neural representations in the hippocampus and lateral prefrontal cortex began as highly aligned across participants, then grew more idiosyncratic to each perceiver over time. Similar effects were observed in the ventral medial prefrontal cortex, but only in one cohort of participants. Regions often linked to social cognition (temporoparietal junction, superior temporal sulcus) evinced strong encoding for group normative information at both timepoints. Taken together, these findings suggest that neural representations of familiar others may initially be predominantly driven by information that is consistent across perceivers (e.g., physical appearance, face-based evaluations, and trait impressions), then become more unique over time as each perceiver gains their own experiences with and impressions of these individuals.

**Significance Statement:** Understanding how humans form mental representations of others is fundamental to the study of social behavior. Yet we know little about how these representations evolve as people get to know each other over time. Using brain imaging in a real-world social network of first-year high school students, we tracked how participants’ neural representations of their classmates changed over the course of a year. While some brain regions encoded stable, shared representations of peers, others showed increasingly individualized patterns. These findings shed light on how real-world social experiences shape representations of familiar others and delineate how the brain balances both personal experience and shared social information. This work informs theories of social cognition by showing how mental representations evolve over time.

## Introduction

Humans live their lives embedded in complex and dynamic social environments. To act adaptively in these contexts, they must encode, store, and use knowledge about the people around them^1^. Doing so involves constructing mental representations of others—cognitive schemas built upon autobiographical experiences and person-specific knowledge, and informed by broader sociocultural scripts^2^. While both theoretical and empirical work has confirmed the importance of mental representations of others for shaping social behavior across a diverse range of contexts and relationship types^3–5^, the temporal dynamics of mental representations are poorly understood. Next to little is known about how mental representations of other people change over time. Do they become more idiosyncratic with one’s unique lived experience, or do they converge on common information? Although some work has examined short-term changes in representations of unfamiliar individuals within laboratory settings^6–9^, in the real world such representations are informed by troves of experience and information involving familiar others that accumulate over protracted timescales, far longer than what is feasible to manipulate in a lab setting.

On one hand, mental representations of other people may become more *similar* across perceivers over time. Early impressions of others may be shaped by limited or unrepresentative contextual information^10^, and thus may be inaccurate or highly variable across perceivers^11^. As perceivers gain more exposure to a given social target, they are likely to observe regularities in behavior, temperament, affect, and preferences. These shared observations may promote convergence in representations across perceivers over time.

On the other hand, mental representations of other people may become more *idiosyncratic* over time—gradually tailored to each perceiver’s unique experiences with a target. This hypothesis is informed by findings from the social perception literature demonstrating that initial impressions and evaluations show remarkable consistency across perceivers, even at zero acquaintance (e.g., perceivers’ impressions tend to align even if only based on a photograph or short description of another person^6^). This initial alignment across perceivers may stem from commonly available perceptual information (e.g., facial features) interacting with shared cultural heuristics and stereotypes^12^. Over time, however, representations may become more idiosyncratic as perceivers accumulate unique experiences with the target, or form distinct reactions to experiences with the target, even if those experiences are shared among perceivers. Representations may also diverge on the basis of second-hand impressions that are propagated within different peer cliques or crowds^13–15^. These processes would result in individualized, idiosyncratic models of the same person across different perceivers.

It is important to stress that both possibilities are supported by evidence that initial impressions of novel social targets are not fixed, and are subject to change based on subsequently presented information^6,16–18^. In other words, the literature is in agreement insofar that it suggests mental representations *can—*perhaps even should— change over time, proffering multiple possible mechanisms by which change occurs. What remains unknown, however, is precisely *how* mental representations change over significant periods of time.

In the present study, we sought to investigate how neural underpinnings of mental representations of familiar others change with social experiences over time. We followed two cohorts of first-year high school students over the course of a school year (Figure 1). Using functional magnetic resonance imaging (fMRI) and representational similarity analysis (RSA), we sought to quantify the degree to which brain regions encode information that is common across perceivers versus information that is idiosyncratic to each perceiver, and how this changes with time. We specifically chose to investigate the neural underpinnings of mental representational change for two key reasons. First, data derived from measures that only capture conscious aspects of mental representations can be inadequate. This is because much of everyday mental life relies heavily on non-conscious operations, and operations that *do* occur with conscious awareness are often marked by limited or inaccurate introspective access^19,20^. Second, existing cross-sectional fMRI studies show that various aspects of storing, accessing, and forming mental representations are widely distributed across numerous brain regions^3,11,21–25^; thus, it is possible that different patterns of results would be observed in different brain regions.

**Figure 1.**
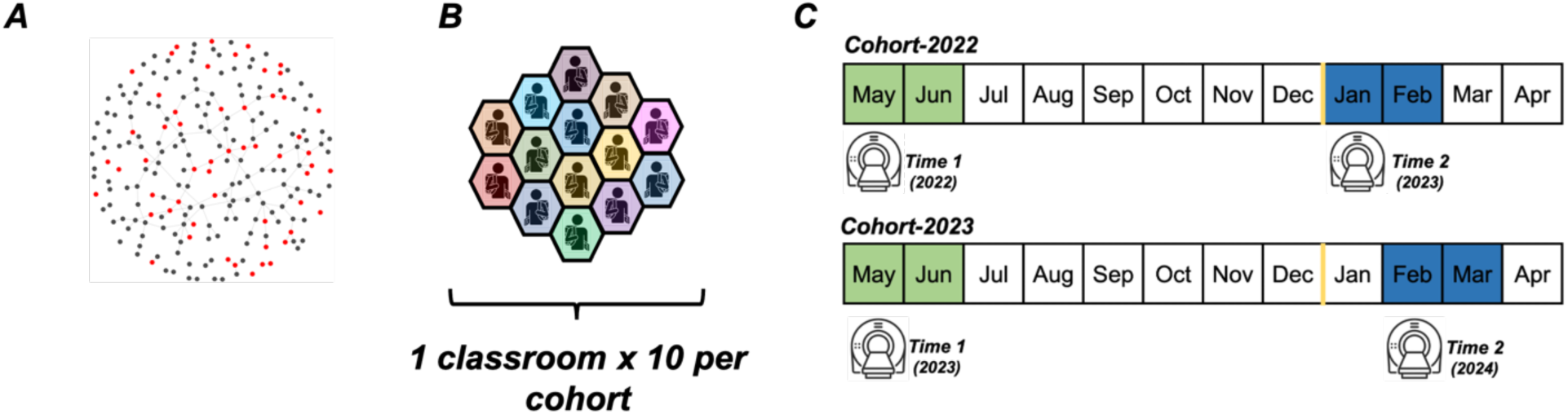
Data collection timeline *Note.* (A) Visualization of one cohort network at single time point, for illustrative purposes (2023 Cohort, Time 1). In each cohort, participants belonged to one of ten classrooms (B) and were scanned twice during the academic year (C). Calendar cells shaded in green and blue represent Time 1 and Time 2 data collection, respectively. The gold line represents the end of the calendar year. The South Korean academic school year begins in March; thus, participants were enrolled into the parent study early on in the school year, with Time 1 scans taking place shortly thereafter.

We were also deliberate in our choice to target a developmentally significant period–i.e., the first year of high school–for this study. Doing so presents a unique opportunity for longitudinal investigation of mental representations of known others for several reasons. First, participants are sorted into a discrete set of peers at the beginning of the academic year (i.e., classrooms), and then receive consistent, daily exposure to these individuals over many months. This allows for the study of how representations of others change over time in a real-world context marked by shared experiences with classmates, frequent interaction, and increasing familiarity. Second, adolescence is a period of heightened social sensitivity and identity exploration. During this time, individuals become neurobiologically oriented towards peer relationships, social dynamics, and are outwardly seeking information from peers for a multitude of reasons^26–28^. Additionally, much of the existing empirical work focuses on representations of people who are not personally known to participants—such as celebrities, confederates, or fictional characters that participants become acquainted with during brief experimental paradigms in temporally constrained, artificial contexts. Of the few studies that do use familiar individuals ^29^, none to our knowledge have examined representational change in a protracted longitudinal study. Testing these hypotheses in an ecologically relevant context that the vast majority of humans experience promises to produce findings that better reflect real-world phenomena.

## Results

### Study & Analytic Overview

Participants (150 adolescent girls) were scanned at the beginning (Time 1) and end (Time 2) of the school year. During scanning, participants completed a face-viewing task where they were shown their classmates’ faces (see Fig. 2 for schematic). RSA was used to quantify the degree to which each participant’s neural representations of their classmates’ faces matched a group norm (i.e., the average of all of their classmates’ neural representations) throughout the brain. We ran tests to uncover brain regions in which individual perceivers’ neural representations of their classmates were aligned with the group norm at each timepoint, and also examined changes in this alignment over time. All analyses were run on an aggregated sample of all participants (statistically adjusting for cohort); however, we also performed *post-hoc* exploratory longitudinal analyses in each cohort separately. To address the growing issue of analytic flexibility in human neuroimaging, all analyses were conducted with multiple specifications and results were aggregated across all of them. Figures of brain-wide statistical maps depict this variability in two ways—one summarizing the density of evidence across specifications, another summarizing the consistency of significant effects (see figure captions). Results reported here come from searchlight analyses; similar results were obtained when re-running analyses iteratively within each parcel of cortical parcellations (see Supporting Information).

**Figure 2.**
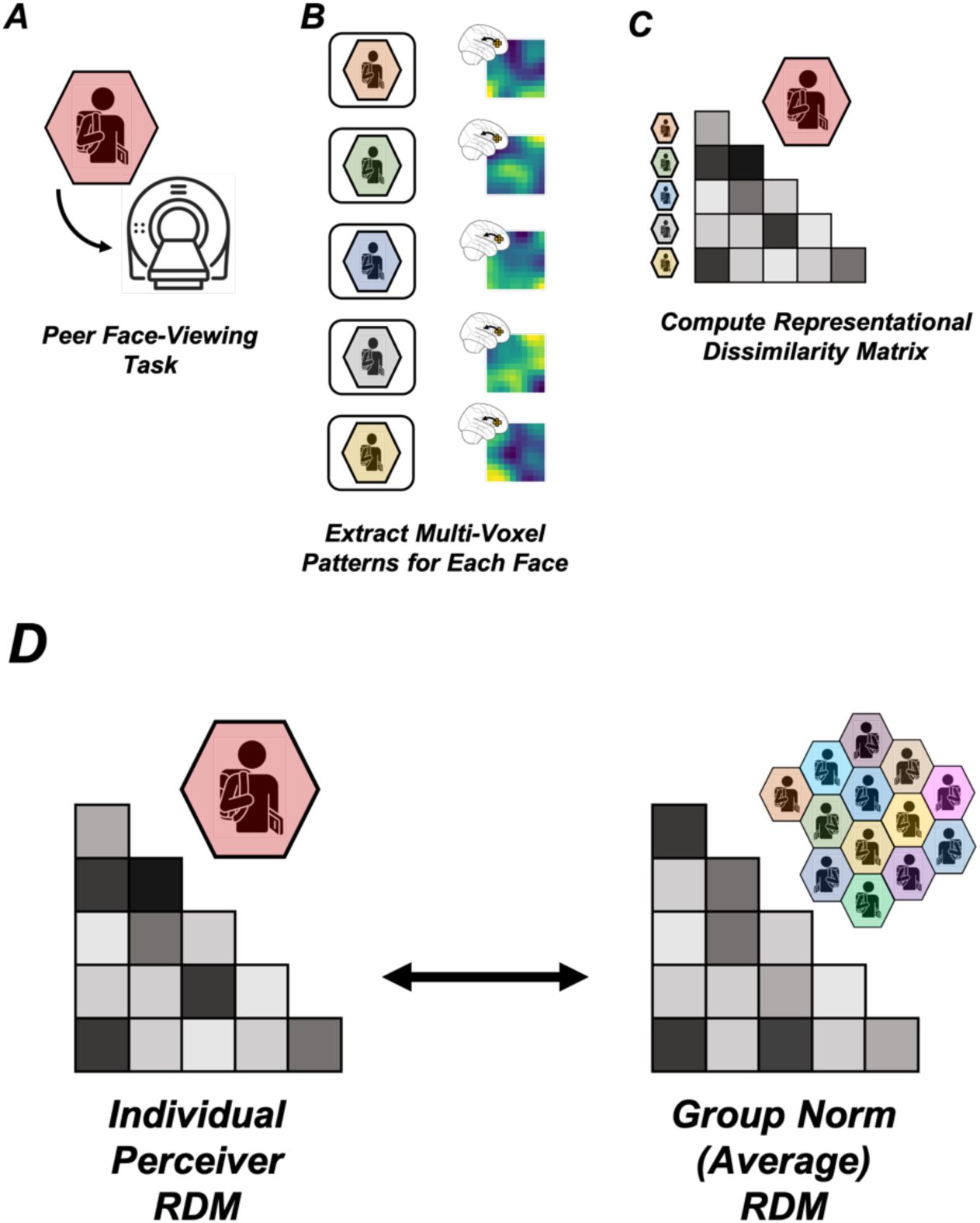
Data analysis schematic *Note.* Participants completed a face-viewing task in which they passively viewed images of their classmates’ faces while undergoing fMRI (A). Each participant’s local multi-voxel response patterns for each face that they viewed were estimated at each point in the brain using a searchlight analysis (B). At each searchlight center, response patterns evoked by each face were compared with one another to generate a representational dissimilarity matrix (RDM; C). Next, at each searchlight center, each participant’s local neural RDM was compared to the group norm (computed by averaging the local neural RDMs from all *other* participants who had seen the same faces; D). Here the perceiver is indicated via red shading, but we emphasize that analyses were iterative (i.e., this process was repeated for each participant in the dataset).

### Several Brain Regions Showed High Representational Similarity Between Individual Perceivers and the Group Norm at the Start and End of the School Year

Figures 3 and 4 depict results of the searchlight analyses probing representational similarity between individual perceivers and the group norm (computed by averaging the local representational similarity matrices (RDMs) of other perceivers, excluding the focal subject; see Methods) at Time 1 and Time 2, respectively.

**Figure 3.**
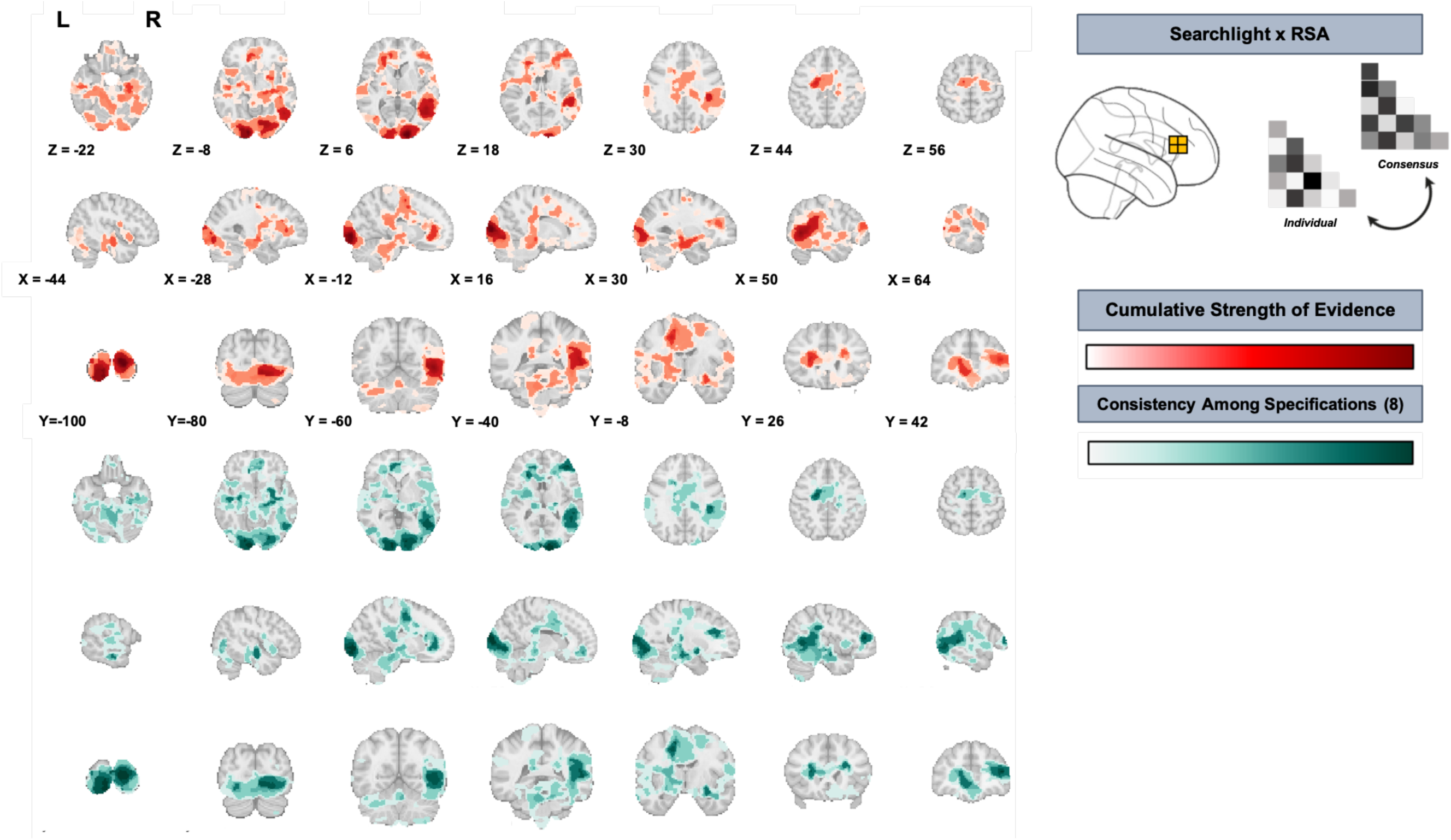
Brain regions where representations of peers showed evidence of robust alignment across perceivers at Time 1. *Note.* The top panel (A) shows the *cumulative strength of evidence* for similarity among participants’ local neural representations of their classmates, computed by summing negative-log10 *p*-values of thresholded and corrected cluster maps across eight different analytic specifications (distance-defining metric: Pearson vs. Euclidean distance, cluster thresholding and correction method: Permutation test vs. max size vs. max mass vs. TFCE). More intense color (red) reflects stronger cumulative evidence across specifications. The bottom panel (B) shows *consistency across specifications*, operationalized as the number of specifications (out of 8 total) in which a given voxel survived statistical thresholding. More intense color (teal) indicates high cross-specification robustness. Results summarized here were obtained with the searchlight method (see Methods).

**Figure 4.**
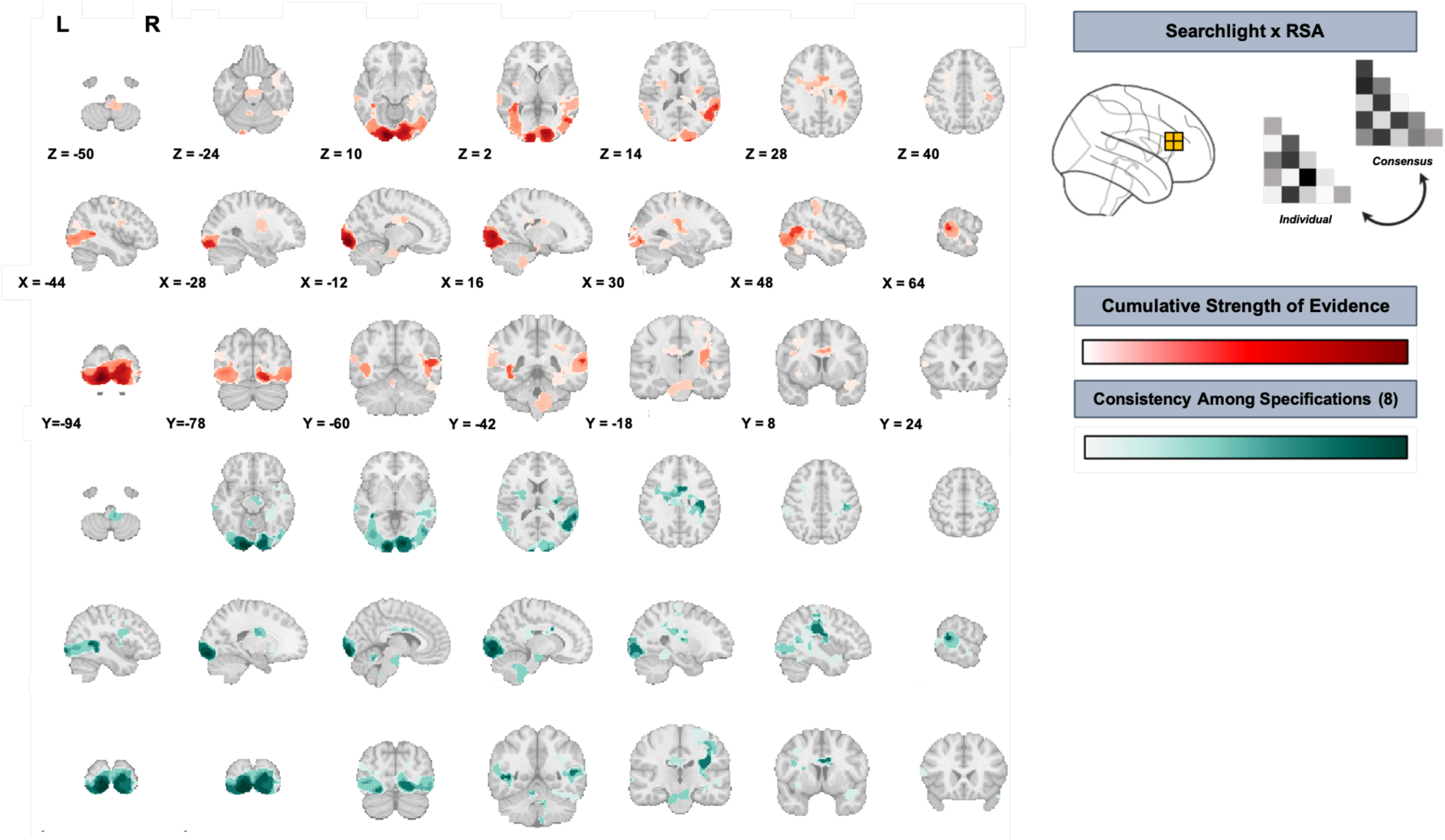
Brain regions where representations of peers showed evidence of robust alignment across perceivers at Time 2. *Note.* The top panel (A) shows the *cumulative strength of evidence* for similarity among participants’ local neural representations of their classmates, computed by summing negative-log10 p-values of thresholded and corrected cluster maps across eight different analytic specifications (distance-defining metric: Pearson vs. Euclidean distance, cluster thresholding and correction method: Permutation test vs. max size vs. max mass vs. TFCE). More intense color (red) reflects stronger cumulative evidence across specifications. The bottom panel (B) shows *consistency across specifications*, operationalized as the number of specifications (out of 8 total) in which a given voxel survived statistical thresholding. More intense color (teal) indicates high cross-specification robustness. Results summarized here were obtained with the searchlight method (see Methods).

At the beginning of the school year (Time 1), we observed significant clusters in areas that include regions involved in visual processing (e.g., lateral and medial occipital cortex; ventral temporal cortex). We also observed significant clusters in the medial prefrontal cortex (mPFC), superior temporal sulcus (STS), hippocampus, parahippocampal cortex, cingulate cortex, and posterior parietal cortex (PPC), including the temporoparietal junction (TPJ); see Fig. 3.

Several months later, at Time 2, we again observed significant alignment of perceivers’ representations in regions involved in visual processing (e.g., occipital cortex), as well as in regions linked to social cognition and memory, including the STS, TPJ, posterior hippocampus, parahippocampal gyrus, cingulate cortex, and anterior temporal cortex (see Fig. 4).

The TPJ, STS, hippocampal, cingulate cortex and occipital lobe findings were present across both timepoints, with the occipital lobe findings being the most robust (Supplementary Figure 1).

### Neural Representations Became More Perceiver-Specific Over Time

Figure 5 depicts results of the searchlight analysis testing for changes in representational similarity between individual perceivers and the group norm across time. Overall, we observed significant clusters in the dorsolateral prefrontal cortex (dlPFC) and the hippocampus, two brain regions implicated in social navigation and higher-order person knowledge ^1,30–32^. In both of these brain regions, neural representations of classmates’ faces became increasingly perceiver-specific over time as participants acquired more experience with one another.

**Figure 5.**
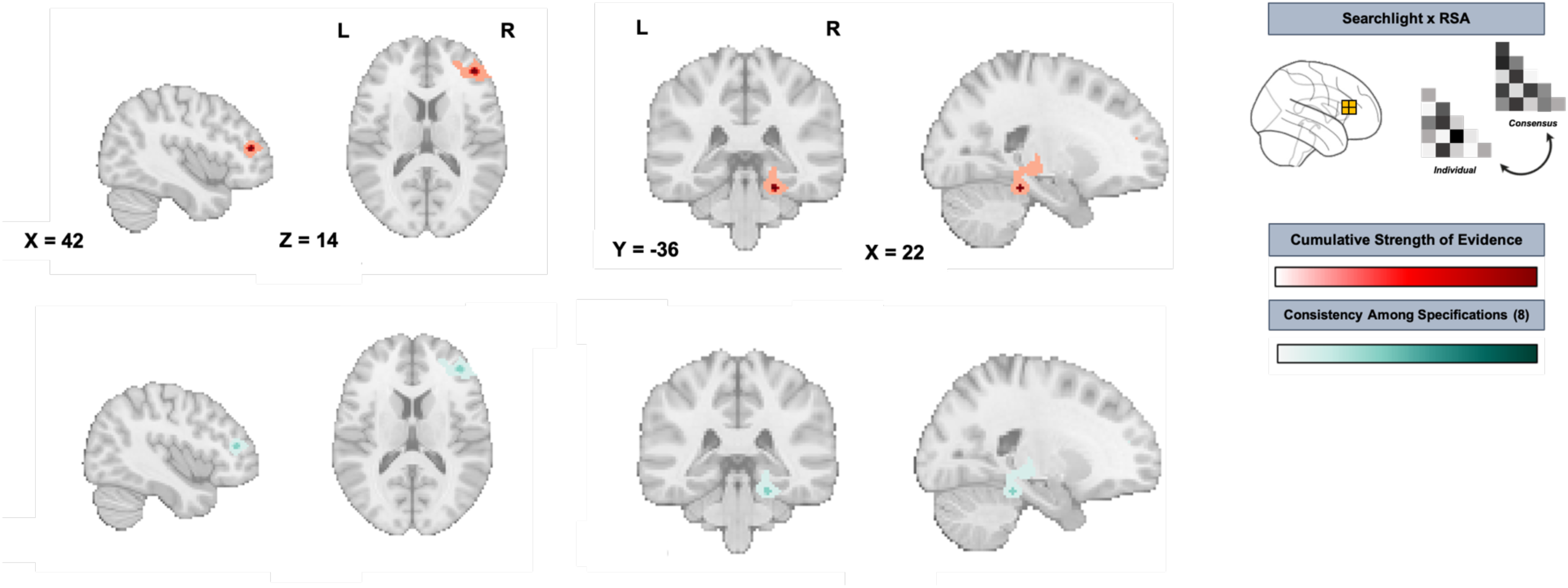
Neural representations became more idiosyncratic to each perceiver over time in the dlPFC and hippocampus *Note.* The **top panel** shows the *cumulative strength of evidence* for representational similarity, computed by summing negative-log10 p-values of thresholded and corrected cluster maps across eight different analytic specifications (distance-defining metric: Pearson vs. Euclidean distance, cluster thresholding and correction method: Permutation test vs. max size vs. max mass vs. TFCE). More intense color (red) reflects stronger cumulative evidence across specifications. The **bottom panel** shows *consistency across specifications*, operationalized as the number of specifications (out of 8 total) in which a given voxel survived statistical thresholding. More intense color (teal) indicates high cross-specification robustness. Results summarized here were obtained with the searchlight method. dlPFC refers to dorsolateral prefrontal cortex.

### Exploratory Longitudinal Analyses Within Each Cohort

We complemented our primary analyses, which involved all participants, with *post-hoc* exploratory analyses that were conducted separately within each cohort. This analysis revealed that neural representations of classmates’ faces also became more idiosyncratic to each perceiver over time in the ventromedial prefrontal cortex (vmPFC), a key hub for social cognitive processing^33^. However, this effect was only observed in the 2023 cohort (Supplementary Figure 2).

When looking within each cohort separately, neural representations became less perceiver-specific over time in portions of the left PPC, although the areas of the PPC that were implicated did not overlap across cohorts (Supplementary Figure 3). We interpret these cohort-specific results with caution, given that this pattern of results did not replicate across cohorts or emerge when aggregating data across cohorts.

## Discussion

The present study leveraged fMRI and a longitudinal design in a sample of first-year high school students to examine how mental representations of familiar individuals change as individuals become better acquainted with each other over time. Guided by competing hypotheses, we tested whether neural representations of others converge across perceivers or diverge to become more idiosyncratic and perceiver-specific. The resultant findings provide rare evidence of how neural representations of personally familiar others evolve following months of real-world interaction. By moving beyond short-term lab-based paradigms that feature artificial or unfamiliar perceptual targets (e.g., fictive individuals, celebrities), these findings offer an ecologically valid window into such phenomena. Across two cohorts, we found that months of rich, varied everyday encounters drive increasingly individualized, perceiver-specific representations in some brain regions (e.g., dlPFC, hippocampus), while also preserving stable, shared aspects of person knowledge and of perceivers’ responses to peers in other brain regions (e.g., occipital cortex, TPJ, STS). Taken together, these results shed light on how real-world experiences shape and personalize mental models of the social world.

Representations in the dlPFC and hippocampus became increasingly perceiver- specific over the course of the school year. Past research has shown that the hippocampus is involved in encoding person knowledge, social navigation, and episodic memory^21,22,34,35^, while the dlPFC is involved in processing social information in contexts requiring deliberative judgments^36–38^. The fact that participants’ representations of their classmates grew more idiosyncratic over time in these regions could reflect integration of unique experiences, interpretations, and evaluations into their mental models of others, coloring such representations with increasing personalization. This suggests that the accumulation of episodic memories of specific interactions and experiences might partly underlie this representational divergence in the hippocampus whereas divergence in the dlPFC may reflect the integration of this information into social judgments.

Exploratory within-cohort analyses found analogous results in the vmPFC in one cohort, suggesting the possibility that valuation or affect-related responses to familiar others also become more idiosyncratic to each perceiver over time, potentially reflecting, for example, the emergence of each perceiver’s unique preferences and/or affiliative or acrimonious relationships with their classmates; however, these vmPFC findings emerged only in one cohort and thus require replication. One potential explanation for the observed vmPFC differences between cohorts relates to COVID-19 mask-mandates and their potential effects on participants’ observations of and interactions with one another in everyday life. Indoor mask usage was mandated in South Korean schools until January 2023. This means that the 2022 Cohort was mandated to wear masks indoors from the beginning of their time together as classmates until the onset of Time 2 data collection (Figure 1), whereas the 2023 Cohort was not. This difference between the cohorts’ social perceptual environments could have impacted the evolution (and in particular, the individualization) of participants’ representations of one another in several ways, including by restricting access to rich facial cues, leading to less person-specific input when participants in the 2022 Cohort interacted with and observed each other between Time 1 and Time 2.

While representations in certain brain regions shifted to become more perceiver- specific over time, we also observed strong alignment of participants’ neural representations of their classmates in several brain regions at the start and end of the year. This included regions often implicated in social cognition and person knowledge, including the STS and TPJ^21,22,34,39^, as well as regions of the occipital lobe that support visual processing ^40,41^. Additionally, although participants’ representations of their classmates became increasingly perceiver-specific over time in the hippocampus, there was still strong alignment across perceivers at both timepoints. These results suggest that aspects of participants’ representations of their classmates that are subserved by these brain regions retained shared information across both perceivers and time; this was the case even in regions where representations grew relatively more idiosyncratic over time (e.g., the hippocampus). This stability could be driven by information that is readily observable to and broadly agreed upon across perceivers and that would remain relatively consistent over time (e.g., facial features apparent in photographs, culturally shared interpretations of such features, enduring traits, behavioral regularities).

These findings provide insight into the protracted process by which neural representations of the people in one’s life evolve over time and become refined by real-world experience. More specifically, this set of findings provides evidence for both the persistence of stable person representations that are shared across perceivers, as well as the gradual emergence of more idiosyncratic representations that contain more perceiver-specific detail. This pattern of results is consistent with prior research on social perception, which has shown that early impressions of other people tend to be driven by shared perceptual cues and cultural heuristics^10^. Our results suggest that following these early impressions, sustained direct interactions as well as network level processes (e.g., secondhand impressions spreading within communities of a network) likely lead individual perceivers to develop more individualized representations of the people around them.

Our results also contain key insights for developmental science, given the adolescent population studied here. Adolescence is a period marked by heightened sensitivity to peer relationships, with teens increasingly attuned to social peer dynamics in an effort to seek autonomous social affiliation, novelty, and content for novel identities^26,42–44^. While lay perceptions of adolescence often depict teens as almost completely reliant on peers for shaping their attitudes, opinions, and identity, the current findings could be seen as at odds with such depictions. One developmental interpretation of these findings is that the adolescent brain does not overwhelmingly default to shared, socially normative representations of others, but instead carries the capacity to develop individualized models based on unique social experiences. In this way, our findings counter the idea that adolescence is purely a period of conformity or susceptibility to peer influence; rather, they point to a nuanced interplay between social affiliation and personal meaning-making. At the same time, the fact that many brain regions contained representations of classmates that were highly aligned across perceivers—particularly early in the school year—is consistent with the relatively rapid uptake of normative social information, reinforcing the importance of peers during this developmental window. Together, these findings provide a more textured account of adolescent social development, in which mental representations evolve through both collective socialization and individual experiential refinement.

Relatedly, it is worth noting that developmental context of this study—an adolescent sample during the first year of high school—likely provides a particularly sensitive context for observing the phenomena studied here. As mentioned above, during adolescence, social sensitivity tends to be heightened and individuals are particularly attuned to their peers^26–28^. Moreover, the school environment creates dense networks of regular and significant social encounters between classmates. As such, the current study’s setting may have provided a particularly sensitive testbed for observing how repeated shared experiences, combined with person-specific interaction histories, shape neural responses to familiar others. Future work should test if similar results are found across diverse settings, age groups, and cultures.

Exploratory *post-hoc* analyses revealed potential cohort-specific effects, including a shift toward idiosyncratic representations in ventromedial prefrontal cortex (vmPFC) and a shift toward shared representations in posterior parietal cortex (PPC). We are cautious in interpreting these cohort-specific effects, since they were not observed in the aggregate sample. Extending the current paradigm to a diversity of samples and settings would also provide an opportunity to examine if the cohort-specific effects observed here would replicate. Future work could also extend on the current findings by sampling participants’ neural representations of each other at more than two time points. While the two-timepoint design used here captures broad changes that occurred over the course of the school year, denser temporal sampling could reveal finer-grained temporal trajectories of representational change as found in recent work in other domains^45,46^. In addition, it would be helpful for future research to incorporate additional behavioral measures to examine what aspects of people’s perceptions, knowledge, memories, and evaluations of their peers underlie the observed neural effects.

In conclusion, the current study demonstrates how neural representations of real-world social partners evolve over months of sustained interactions. Neural representations showed significant alignment across time and perceivers, while also becoming increasingly individualized over time through perceivers’ lived experiences.

By examining these processes in an ecologically valid setting and over a protracted time scale, these results provide insight into how the brain constructs and personalizes representations of the social world.

## Methods

### Participants

Participants from the current report were drawn from a broader, multi-cohort longitudinal study investigating socioemotional development in adolescence. The sample here comprises a *N* = 150 subset of first-year high school students from an all-girls school in Seoul, South Korea who completed an fMRI component. The sample of fMRI participants here consists of individuals from one of two cohorts of students that either began high school in 2022 (*N_2022_* = 70, mean age = 15.88 years, SD = 0.31) or in 2023 (*N_2023_* = 79, mean age = 15.89, SD = 0.31). Because the broader study is an ongoing sociological investigation of a single school, sample size was determined by the number of students within the school who would participate. Participants belonged to one of 10 individual classrooms per cohort, totaling 20 individual unique classrooms across the entire sample.

All participants participated at least one timepoint. Any cross-sectional analyses include all participants that provided data for the relevant time point; longitudinal analyses comprise participants that participated at each time point. For the 2022 Cohort, 43 participants had available data at both time points, 14 participants only had available data at Time 1, and 14 additional participants only had available data at Time 2. For the 2023 Cohort, 48 participants had available data at both timepoints, 16 participants only had available data at Time 1, and an additional 15 participants only had available data at Time 2. Participants were compensated ₩100,000 (equal to approximately $70-$80 USD at the time of data collection) for completing the MRI scan. The research was completed at Yonsei University. Written assent and consent were obtained from all participants and their parents, respectively. All research protocols were approved by the Institutional Review Board at Yonsei University.

### Experimental Protocol

Participants from both cohorts were scanned twice, once at the beginning of the academic term (Time 1) and another at the end (Time 2). Scans for Time 1 for the 2022 and 2023 cohorts took place between May and June of 2022 and 2023, respectively.

Scans for Time 2 took place between January-February and February-March of the following year for the 2022 and 2023 cohorts, respectively (see Fig. 1).

### MRI Acquisition Parameters

Functional neuroimaging data for the 2022 cohort were collected using a 3T Siemens Magnetom Trio scanner and a 32-channel head coil located at Seoul National University. A high resolution T1 magnetization-prepared rapid-acquisition gradient echo (MPRAGE) structural image was acquired for registration purposes (TR = 2300 ms, TE = 2.36 ms, Flip Angle = 9°, FOV = 256 mm^2^, 1 mm^3^ isotropic voxels, 224 slices, A >> P phase encoding). Functional runs were comprised of T2*-weighted echoplanar images (TR = 2000 ms, TE = 25 ms, Flip Angle = 80°, FOV = 208 mm^2^, 3 mm^3^ isotropic voxels, 38 slices, A >> P phase encoding).

Imaging data for the 2023 cohort were collected using a 3T Siemens Vida scanner scanner and 64-channel head coil located at Yonsei University. A high resolution T1 MPRAGE scan was collected (TR = 2300 ms, TE = 2.26 ms, Flip Angle = 8°, FOV = 256 mm^2^, 1 mm^3^ isotropic voxels, 192 slices, A >> P phase encoding) in addition to T2*-weighted echoplanar functional runs (TR = 1160 ms, TE = 30 ms, Flip Angle = 63°, FOV = 208 mm^2^, 3 mm^3^ isotropic voxels, 56 slices, A >> P phase encoding). Scan parameters were consistent for both timepoints within cohort.

### Face-Viewing Task

All participants completed two runs of a face-viewing task in the scanner using a rapid event-related design. Each run consisted of 42 trials where participants viewed the faces of six of their classmates. The six classmates were chosen based on peer popularity ratings that were collected as an additional measure from the broader study (two popular, two unpopular, two neutral). This measure was used for separate analyses targeting a different set of research questions and thus is not further discussed here. Participants were only shown faces from their own classrooms. Within a given classroom, participants saw overlapping but slightly different sets of faces, which means that the group norm RDM differed slightly for each participant. On average, each face presented to participants at a given timepoint was also viewed by approximately 4–6 *other* participants in the same classroom within the same cohort (2022 Cohort: an average of 4.93 other participants at Time 1, 4.61 at Time 2; 2023 Cohort: 5.46 at Time 1, 6.18 at Time 2). Each face was shown between six to nine times on each run (totaling 12-18 total presentations per face across the two runs). Faces were presented for 2,000 ms followed by a jittered intertrial-interval lasting between 4,000 – 11,000 ms (average = 5,500 ms). Indoor mask-wearing in South Korean schools was mandated due to COVID-19 until January 2023. Thus, all participants in the 2022 Cohort would have been masked when interacting with and observing one another indoors until the onset of Time 2 data collection. While mask-wearing was not mandated for the 2023 Cohort, it was not unusual for participants to wear masks in the classroom during this time period. Therefore, half of the images in the face-viewing task depicted each classmate wearing a mask and half did not. Unmasked stimuli depicted classmates with a neutral facial expression in a front-facing ‘headshot’ format. Each run lasted approximately 5.5 minutes.

Complete and systematic data about whether participants were personally acquainted with their classmates prior to starting high school were not collected. However, we did obtain information about which middle schools participants had attended before starting high school, and thus, which participants had attended the same middle schools as each other (although we note that participants would not necessarily be familiar with all students from their middle school). The majority (63%) of participants only saw the faces of other students who had not attended the same middle school as them. On average, approximately 92% (5.5/6) of the people in each participant’s stimulus set were classmates who had attended a different middle school than their own.

### Preprocessing and Statistical Analysis

*fMRI Data Preprocessing.* Preprocessing was performed using fMRIPrep^47^ 20.2.5, which is based on Nipype 1.6.1^48,49^. Boilerplate preprocessing details can be accessed in the Supporting Information.

### Multi-Voxel Pattern Estimation

Multi-voxel patterns were estimated by modeling imaging data from the face-viewing task using a General Linear Model (GLM). Both runs for each subject’s data were modeled using a fixed effects GLM in the NiLearn python package. Each of the 6 faces that each participant saw in the scanner were entered as their own regressor. GLM trial durations were fixed to match the task timing (2000 ms). These regressors were convolved with the hemodynamic response function (double gamma)^50^. Temporal derivatives were added to account for slice timing effects. Several nuisance regressors were also added to statistically control for noise and head motion: the first five aCompCor components, 24 head-motion regressors (6 translation, 6 rotation, their derivatives, and their squared derivatives), and individual volumes exceeding the FD and DVARS thresholds, all computed by fMRIPrep. Finally, a discrete cosine transform was performed by adding a set of slow oscillating functions to the GLM to serve as a high pass filter (.01 Hz). HRF convolution and the discrete cosine transform were performed using Nilearn’s functionalities within the `make_first_level_design_matrix` module. First-level GLMs were run on the preprocessed bold data in MNI152Nlin2009cAsym space. Linear contrasts comparing each face to baseline were computed. Data were not spatially smoothed to preserve fine-grained spatial detail in multivoxel response patterns. The resultant *t*-statistic map for each contrast was used for multivoxel pattern analyses described in the next section.

### Representational Similarity Analysis

Representational similarity analysis (RSA)^51^ was used to compare the neural representations of classmates across participants (specifically, between each individual participant and the average of their classmates). We conducted RSA using a searchlight method ^52^ in addition to running it iteratively in individual parcels of a whole-brain cortical parcellation. The rationale for doing so was to leverage the complementary strengths of these approaches. The *searchlight* approach can reveal results localized to small patches of cortex that may be obfuscated if only examining entire parcels; it also provides sensitivity to results that do not respect cortical boundaries (e.g., effects within a cube that sits at the border of two parcels). On the other hand, the *parcellation* approach aggregates responses within functionally defined regions whose boundaries are derived from the clustering of intrinsic functional connectivity and constrained by the anatomy of the cortical surface, rather than a shape (e.g., a searchlight cube) that does not correspond to neural anatomy or functional organization. In addition to acting as *de facto* spatial priors that help align findings with known cortical organization, this approach can also increase statistical power by aggregating results into a smaller number of units to analyze. Using both approaches also provides a way to gauge the robustness of our results across different analytic specifications, an increasingly important issue in human neuroimaging^53,54^.

The following procedure was implemented by iterating for each subject in the dataset for both the searchlight and parcel analyses (see Figure 2 for an overview). For each focal subject, an RDM was created by computing pairwise distances between multi-voxel patterns of brain activity elicited by each face viewed in the scanner (extracted from the *t*-statistic contrast images described in the preceding section). Next, we identified all other subjects who saw the same faces as the focal subject and extracted the corresponding multi-voxel patterns based on their *t*-statistics. Separate RDMs were computed for each of these non-focal subjects, and these RDMs were then averaged to generate the group norm RDM to compare to this participant’s RDM at this point in the brain. We used both Pearson’s *r*, which captures similarity in locally distributed response patterns, independent of overall response magnitude, and Euclidean distance, which is sensitive to differences in both overall response magnitude and in the spatial distribution of response magnitudes across voxels, to construct the subject-specific and group-average RDMs; doing so provided complementary perspectives on the representations being characterized^55^. Because participants in a given classroom saw overlapping but slightly different sets of faces, this meant there were unique subject-specific and group norm RDMs for each individual in the dataset in each region (i.e., at each searchlight center and within each parcel) of the brain.

Representational structures were compared by using Pearson’s *r* to correlate the unique off-diagonal elements of the two RDMs (subject-specific, group norm). For the searchlight analysis, the resulting correlation coefficient was saved to each center voxel of the searchlight (2mm radius, 125 voxel cube, ∼905 mm^3^; searchlight masks dilated using the searchlight radius). Searchlight maps for each individual subject were saved and used in group-level analyses (see next section). The searchlight procedure was implemented in the brain imaging analysis kit (brainIAK, http://brainiak.org, RRID:SCR_014824) python software package^56^ (Kumar et al., 2020). For analyses using parcellations, one coefficient was estimated for every parcel. Individual coefficient values per parcel for each subject were saved for group-level analyses. We used the 7 networks, 400 parcel version of the Schaefer parcellation^57^.

### Group-Level Analyses

Two types of group-level analyses were performed. The first type tested for brain regions showing similarity between individual participants’ representations and the average of other perceivers who saw the same faces; this analysis was repeated separately at Time 1 and Time 2. The second type tested whether any brain regions showed significant changes in cross-perceiver representational similarity over the two timepoints. Both types of group-level tests were run on the subject-level searchlight and parcellation maps, albeit with slight differences based on the approach. We statistically adjusted for cohort membership in all analyses.

For the searchlight approach, we first implemented a non-parametric permutation equivalent of a one-sided *t*-test using Nilearn’s `non_parametric_inference` module. We specified 10,000 permuted iterations, applied 5.00 mm smoothing (full-width at half maximum), and set a cluster-defining *p*-value threshold of 0.001 (with one exception below). To make sure our findings were robust to analytic flexibility, we separately employed four different types of correction methods to correct for multiple comparisons and ensure a family wise error rate of 0.05: voxel-level (corrected based on the distribution of maximum test statistics from the permutations), cluster size (spatial extent), cluster mass (spatial extent combined with cluster height) and threshold-free cluster enhancement (Smith & Nichols, 2009; this method used various cluster-defining thresholds). This was performed for both timepoints. An effects coded indicator variable (1 = 2022 cohort, -1 for 2023 cohort) was used to adjust for cohort membership.

Next, we examined paired differences across timepoints with the same permutation test by using a design matrix with a column coding change over time (1 = Time 2, -1 = Time 1), a set of overparameterized dummy-coded variables to adjust for random participant effects (e.g., each column contained all 0 except for two entries of 1 corresponding to the entries in the Y vector for a given subject’s two timepoints), and dummy coded indicator (1 = 2022 cohort, 0 for 2023 cohort) to adjust for cohort membership. The same four cluster correction methods were used. This analysis was repeated twice with different signs on the indicator variable for paired differences to test for both increases and decreases in representational similarity across time.

For the parcellation approach, we first used ordinary least squares regression to test whether each parcel evinced significant representational similarity. A design matrix with a column of 1s and the same aforementioned effects coded indicator variable was used. The coefficient related to the intercept, representing the grand mean of a given parcel’s representational similarity after adjusting for cohort, was tested against zero.

This process was repeated for every parcel, with a *p*-value obtained for each parcel. We corrected for multiple comparisons using two different types of multiple comparison corrections: Holm-Bonferroni to control the family-wise error rate at 0.05, and Benjamini-Hochberg to control the false discovery rate at 0.05. Paired differences were tested at the parcel level using the same design matrix as described for the searchlight approach. The same two multiple comparison correction strategies were again used.

## Author Contributions

JFGM was involved in conceptualization, data curation, data analysis, visualization, and writing (original draft, editing); KS was involved in methodology, data collection, data curation and writing (reviewing & editing); YLS was involved in data curation and writing (reviewing & editing); SS was involved in conceptualization, methodology, and writing (reviewing & editing); YY was involved in conceptualization, funding acquisition, methodology, resources, supervision, and wiring (reviewing & editing); CP was involved in conceptualization, data analysis, supervision, and writing (original draft, editing).

