## Supporting Information, Appendix for "Getting to Know You: Neural Representations of Other People Grow More Perceiver-Specific Over Time"

**Guassi Moreira et al.,**

*Expanded fMRI Data Preprocessing Details.* Preprocessing was performed using fMRIPrep 20.2.5 (Esteban, Markiewicz, et al. 2018; Esteban, Blair, et al. 2018, RRID:SCR_016216), which is based on Nipype 1.6.1 (Gorgolewski et al. 2011; Gorgolewski et al. 2017; RRID:SCR_002502).

The T1-weighted (T1w) image was corrected for intensity non-uniformity (INU) with N4BiasFieldCorrection (Tustison et al. 2010), distributed with ANTs 2.3.3 (Avants et al. 2008, RRID:SCR_004757), and used as T1w-reference throughout the workflow. The T1w-reference was then skull-stripped with a Nipype implementation of the antsBrainExtraction.sh workflow (from ANTs), using OASIS30ANTs as target template. Brain tissue segmentation of cerebrospinal fluid (CSF), white-matter (WM) and gray-matter (GM) was performed on the brain-extracted T1w using fast (FSL 5.0.9, RRID:SCR_002823, Zhang, Brady, and Smith 2001). Brain surfaces were reconstructed using recon-all (FreeSurfer 6.0.1, RRID:SCR_001847, Dale, Fischl, and Sereno 1999), and the brain mask estimated previously was refined with a custom variation of the method to reconcile ANTs-derived and FreeSurfer-derived segmentations of the cortical gray-matter of Mindboggle (RRID:SCR_002438, (Klein et al., 2017). Volume-based spatial normalization to one standard space (MNI152NLin2009cAsym) was performed through nonlinear registration with antsRegistration (ANTs 2.3.3), using brain-extracted versions of both T1w reference and the T1w template. The following template was selected for spatial normalization: ICBM 152 Nonlinear Asymmetrical template version 2009c [Fonov et al. (2009), RRID:SCR_008796; TemplateFlow ID: MNI152NLin2009cAsym].

For each of the 2 BOLD faceviewing task runs per subject, the following preprocessing was performed. First, a reference volume and its skull-stripped version were generated using a custom methodology of fMRIPrep. A B0-nonuniformity map (or fieldmap) was estimated based on a phase-difference map calculated with a dual-echo GRE (gradient-recall echo) sequence, processed with a custom workflow of SDCFlows inspired by the epidewarp.fsl script and further improvements in HCP Pipelines (Glasser et al. 2013). A deformation field to correct for susceptibility distortions was estimated based on fMRIPrep's fieldmap-less approach. The deformation field is that resulting from co-registering the BOLD reference to the same-subject T1w-reference with its intensity inverted (Wang et al. 2017; Huntenburg 2014). Registration is performed with antsRegistration (ANTs 2.3.3), and the process regularized by constraining deformation to be nonzero only along the phase-encoding direction, and modulated with an average fieldmap template (Treiber et al. 2016). The fieldmap was then co-registered to the target EPI (echo-planar imaging) reference run and converted to a displacements field map (amenable to registration tools such as ANTs) with FSL's fugue and other SDCflows tools. Based on the estimated susceptibility distortion, a corrected EPI (echo-planar imaging) reference was calculated for a more accurate co-registration with the anatomical reference. The BOLD reference was then co-registered to the T1w reference using bbregister (FreeSurfer) which implements boundary-based registration (Greve & Fischl, 2009). Co-registration was configured with six degrees of freedom. Head-motion parameters with respect to the BOLD reference (transformation matrices, and six corresponding rotation and translation parameters) are estimated before any spatiotemporal filtering using mcflirt (FSL 5.0.9) (Jenkinson et al., 2002). The BOLD time-series (including slice-timing correction when applied) were resampled onto their original, native space by applying a single, composite transform to correct for head-motion and susceptibility distortions. These resampled BOLD time-series will be referred to as preprocessed BOLD in original space, or just preprocessed BOLD. The BOLD time-series were resampled into standard space, generating a preprocessed BOLD run in MNI152Nlin2009cAsym space. First, a reference volume and its skull-stripped version were generated using a custom methodology of fMRIPrep.

Several confounding time-series were calculated based on the preprocessed BOLD: framewise displacement (FD), DVARS and three region-wise global signals. FD was computed using two formulations following Power (absolute sum of relative motions, (Power et al., 2014)) and Jenkinson (relative root mean square displacement between affines, Jenkinson et al. 2002). FD and DVARS are calculated for each functional run, both using their implementations in Nipype (following the definitions by Power et al. 2014). The three global signals are extracted within the CSF, the WM, and the whole-brain masks. Additionally, a set of physiological regressors were extracted to allow for component-based noise correction (CompCor, Behzadi et al. 2007). Principal components are estimated after high-pass filtering the preprocessed BOLD time-series (using a discrete cosine filter with 128s cut-off) for the two CompCor variants: temporal (tCompCor) and anatomical (aCompCor). tCompCor components are then calculated from the top 2% variable voxels within the brain mask. For aCompCor, three probabilistic masks (CSF, WM and combined CSF+WM) are generated in anatomical space. The implementation differs from that of Behzadi et al. in that instead of eroding the masks by 2 pixels on BOLD space, the aCompCor masks are subtracted a mask of pixels that likely contain a volume fraction of GM. This mask is obtained by dilating a GM mask extracted from the FreeSurfer's aseg segmentation, and it ensures components are not extracted from voxels containing a minimal fraction of GM. Finally, these masks are resampled into BOLD space and binarized by thresholding at 0.99 (as in the original implementation). Components are also calculated separately within the WM and CSF masks. For each CompCor decomposition, the k components with the largest singular values are retained, such that the retained components' time series are sufficient to explain 50 percent of variance across the nuisance mask (CSF, WM, combined, or temporal). The remaining components are dropped from consideration. The head-motion estimates calculated in the correction step were also placed within the corresponding confounds file. The confound time series derived from head motion estimates and global signals were expanded with the inclusion of temporal derivatives and quadratic terms for each (Satterthwaite et al. 2013). Frames that exceeded a threshold of 0.5 mm FD or 1.5 standardised DVARS were annotated as motion outliers. All resamplings can be performed with a single interpolation step by composing all the pertinent transformations (i.e. head-motion transform matrices, susceptibility distortion correction when available, and co-registrations to anatomical and output spaces). Gridded (volumetric) resamplings were performed using antsApplyTransforms (ANTs), configured with Lanczos interpolation to minimize the smoothing effects of other kernels (Lanczos 1964). Non-gridded (surface) resamplings were performed using mri_vol2surf (FreeSurfer).

*Supplementary Figure 1. Brain regions that evinced high representational similarity between the group norm and individual perceivers at both Time 1 and Time 2*

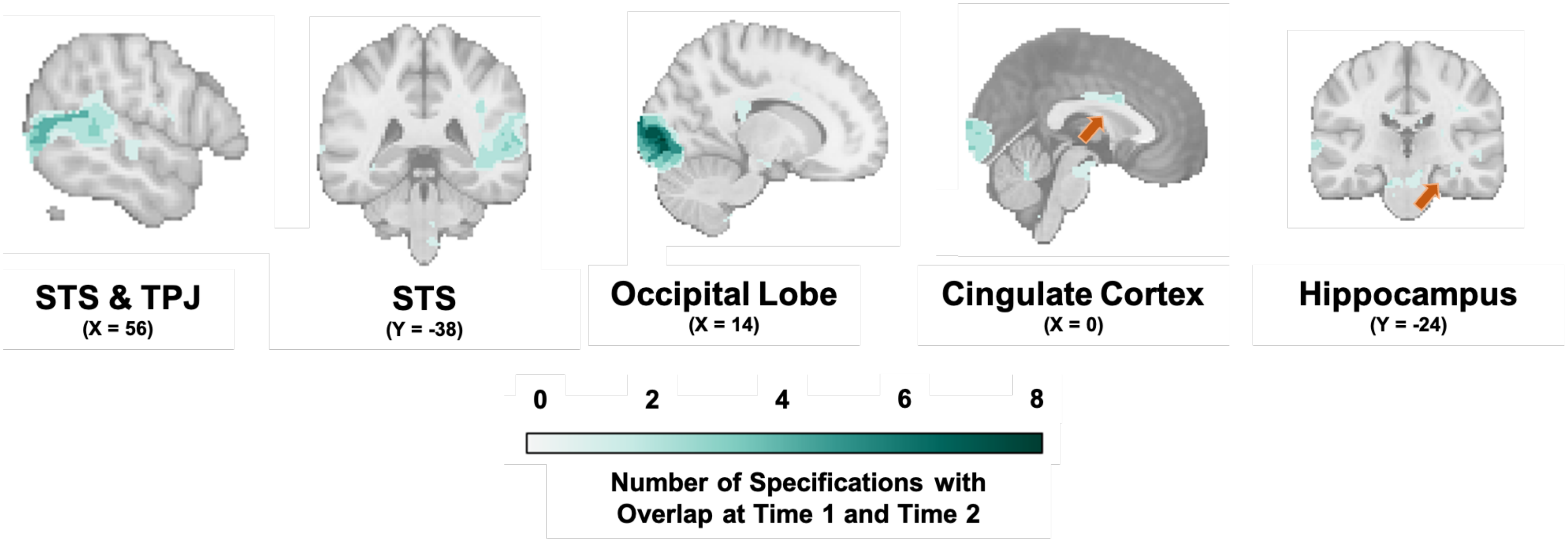


*Note.* STS refers to superior temporal sulcus; TPJ refers to temporoparietal junction. Data for this figure were obtained by first taking the intersection of Time and Time 2 significant voxels for a given specification and then summing the number of specifications with overlap at each voxel.

*Supplementary Figure 2*. Neural representations became more idiosyncratic to each perceiver over time in vmPFC in the 2023 cohort.

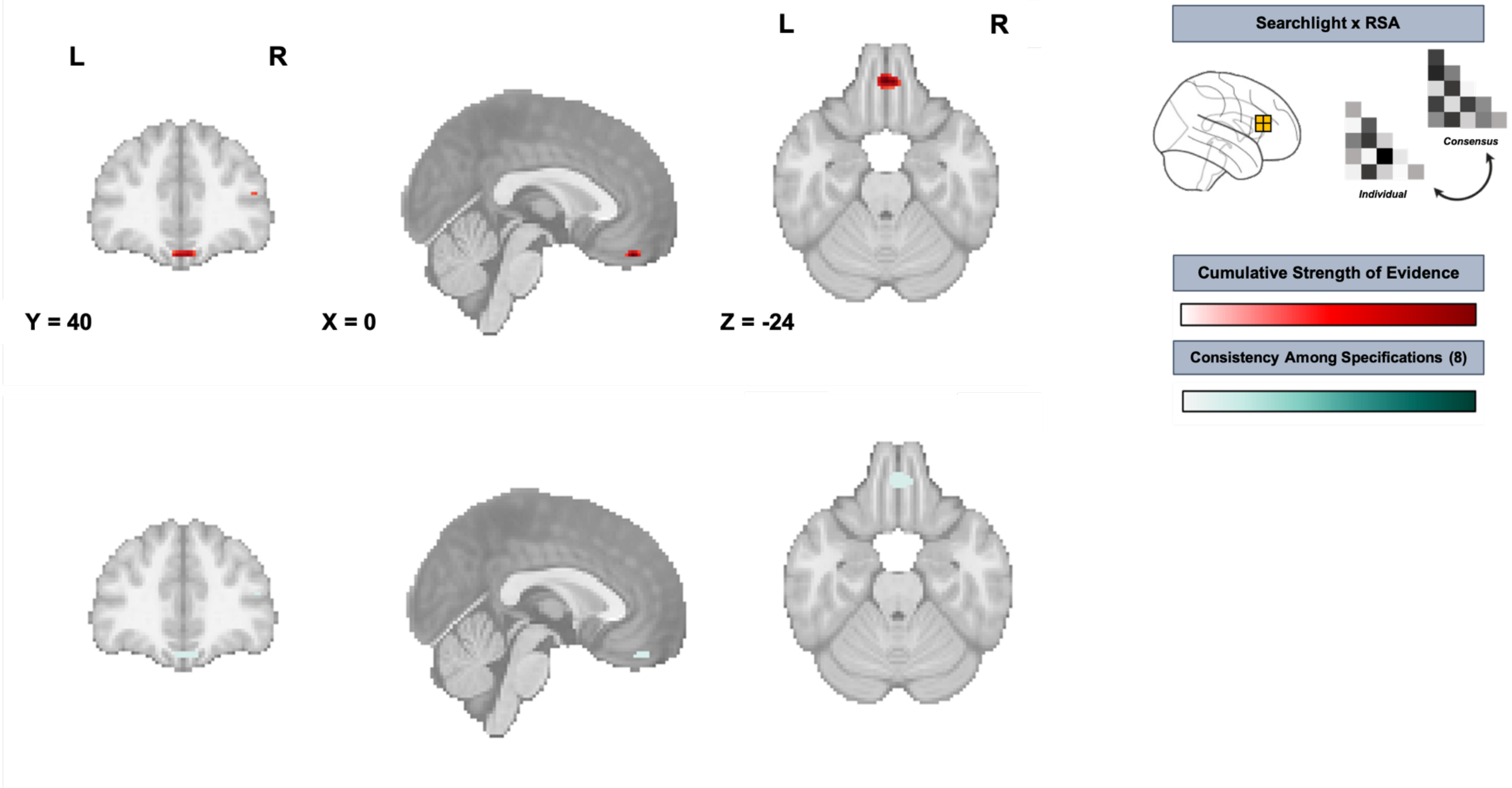


*Note.* The **top panel** shows the *cumulative strength of evidence* for representational similarity, computed by summing negative-log10 p-values of thresholded and corrected cluster maps across eight different analytic specifications (distance-defining metric: Pearson vs. Euclidean distance, cluster thresholding and correction method: Permutation test vs. max size vs. max mass vs. TFCE). More intense color (red) reflects stronger cumulative evidence across specifications. The **bottom panel** shows *consistency across specifications*, operationalized as the number of specifications (out of 8 total) in which a given voxel survived statistical thresholding. More intense color (teal) indicates high cross-specification robustness. Results summarized here were obtained with the searchlight method. vmPFC refers to ventromedial prefrontal cortex.

*Supplementary Figure 3.* Neural representations became more aligned across perceivers over time in different regions of the PPC in each cohort.

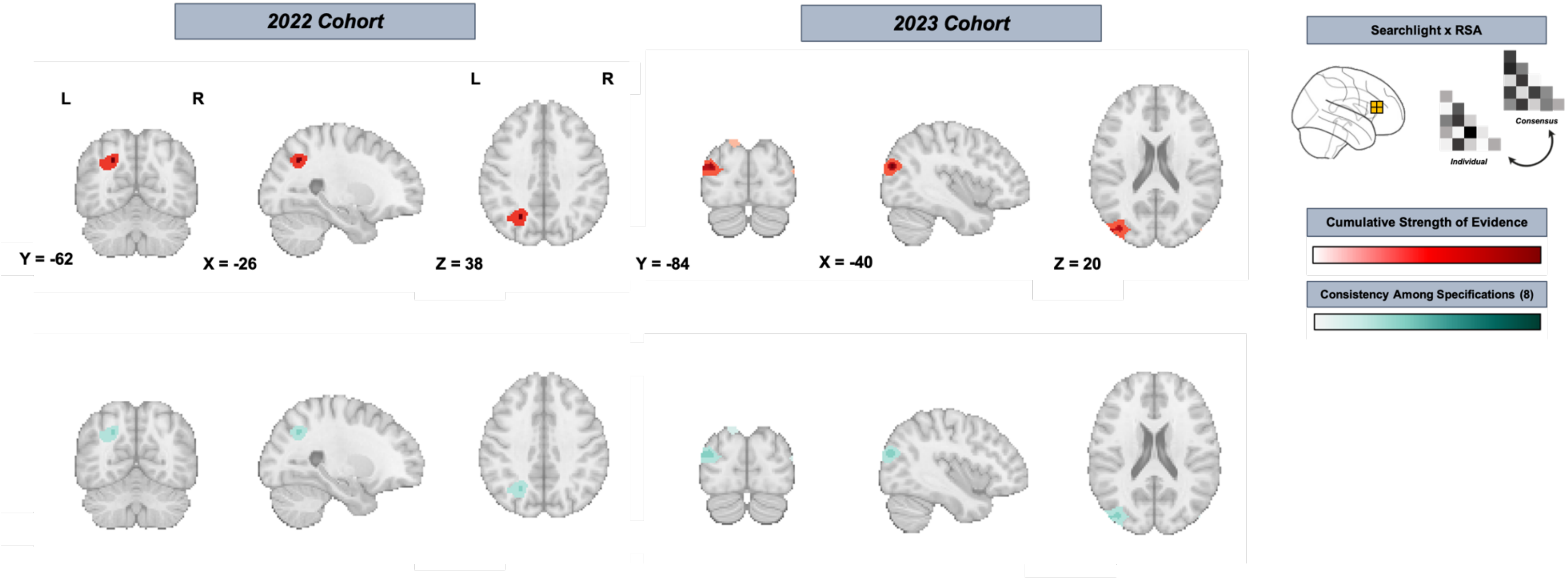


*Note.* The **top panel** shows the *cumulative strength of evidence* for representational similarity, computed by summing negative-log10 p-values of thresholded and corrected cluster maps across eight different analytic specifications (distance-defining metric: Pearson vs. Euclidean distance, cluster thresholding and correction method: Permutation test vs. max size vs. max mass vs. TFCE). More intense color (red) reflects stronger cumulative evidence across specifications. The **bottom panel** shows *consistency across specifications*, operationalized as the number of specifications (out of 8 total) in which a given voxel survived statistical thresholding. More intense color (teal) indicates high cross-specification robustness. Results summarized here were obtained with the searchlight method. PPC refers to parietal posterior cortex. Note that these results should be interpreted with some caution, as these effects did not replicate across cohorts.

*Supplementary Figure 4.* Neural representations became more idiosyncratic to each perceiver over time in the dlPFC (parcellation-based analyses)

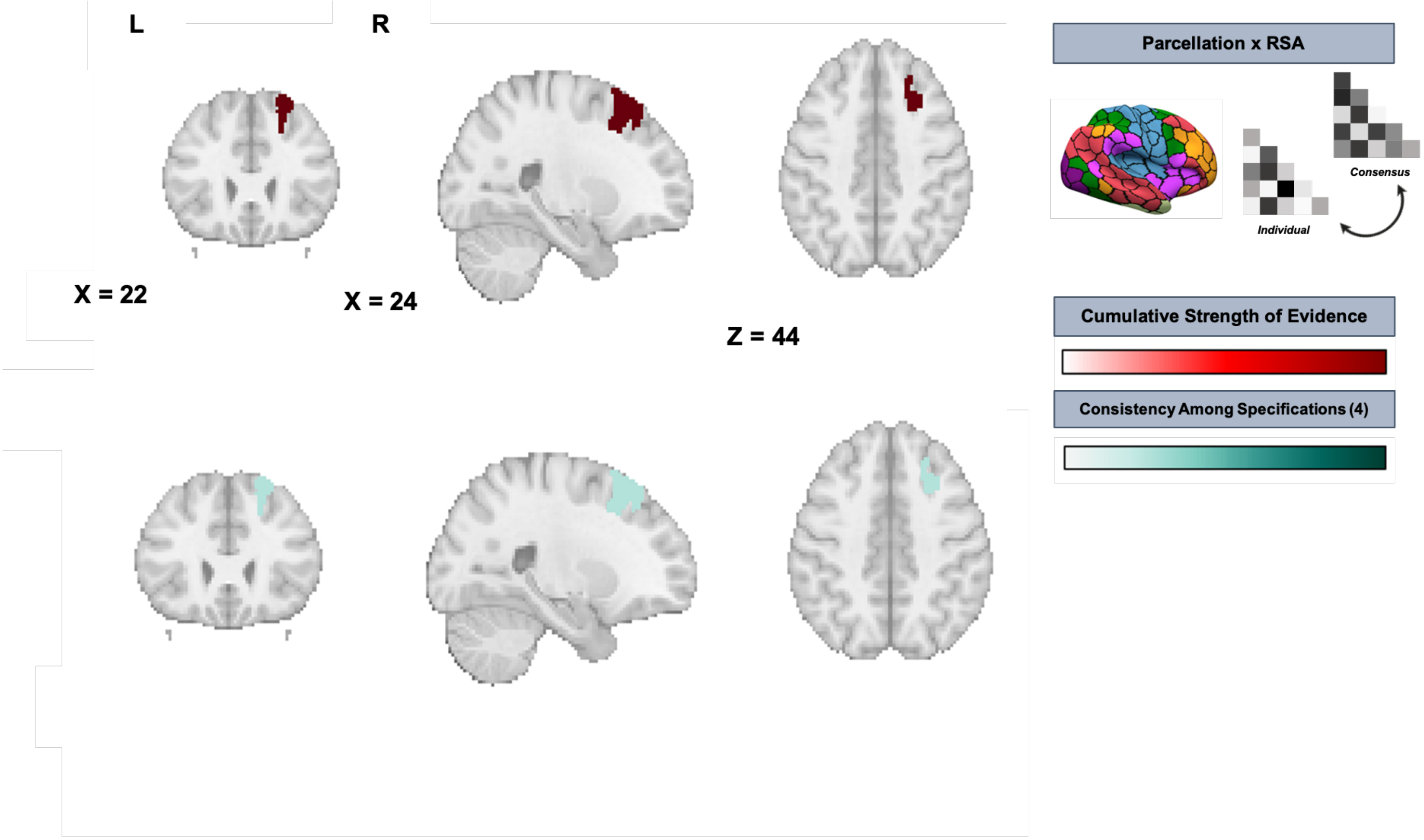


*Note.* The **top panel** shows the *cumulative strength of evidence* for representational similarity, computed by summing negative-log10 p-values of thresholded and corrected cluster maps across eight different analytic specifications (distance-defining metric: Pearson vs. Euclidean distance, multiple comparison correction method: False discovery rate vs. familywise error rate). More intense color (red) reflects stronger cumulative evidence across specifications. The **bottom panel** shows *consistency across specifications*, operationalized as the number of specifications (out of 8 total) in which a given voxel survived statistical thresholding. More intense color (teal) indicates high cross-specification robustness. Results summarized here were obtained on parcellated data. dlPFC refers to dorsolateral prefrontal cortex.
